# Evaluating the differential expression of TAM family receptors and efferocytosis activities in differentiated and polarized THP-1 macrophage

**DOI:** 10.1101/2022.06.21.497076

**Authors:** Megan Chamberland, Brian Farrell, Johannes Yeh

## Abstract

Tissue homeostasis is tightly balanced between cell death and renewal. Each day, as many as 10^11^ cells die in the human body that need to be removed and replaced. The clearance of apoptotic cells, termed efferocytosis, is crucial to tissue homeostasis. Central to this process is macrophage-mediated efferocytosis. Apart from general phagocytosis, efferocytosis is apoptotic cell-specific, which aids in clearing dead cells and preventing the accumulation of self-antigens released by the apoptotic cells. The TAM family receptor kinases (TYRO3, AXL, MERTK) and ligands (GAS6 and PROS) play important roles in engaging the apoptotic cells to initiate efferocytic engulfment and the downstream cellular responses. Dysregulated efferocytic function in macrophages is associated with human diseases such as atherosclerosis, lupus, lung fibrosis, and cancer. Conversely, understanding the regulation and molecular mechanisms of macrophage efferocytosis can potentially lead to beneficial treatments for the above diseases. Despite numerous efferocytosis studies that use primary and cell line-derived macrophages, there has not been a thorough characterization of a cell line system that can be reliably used for efferocytosis assays. Consequently, many macrophage efferocytosis assays reported do not clearly distinguish efferocytosis from phagocytosis. Here we evaluated the THP-1 cell line as a potential human macrophage cell line system for efferocytosis studies. Consequently, many macrophage efferocytosis assays reported do not clearly distinguish efferocytosis from phagocytosis. Through the study we examined the differential expression of the TAM family receptors and their ligands in the various THP-1 macrophage differentiation and polarization states. We also characterized the THP-1 cell line as a reliable system for performing *in vitro* efferocytosis studies.

## Introduction

The process of cell death and renewal is an innate cellular mechanism essential to the maintenance of tissue homeostasis. The apoptotic cell removal process, termed efferocytosis, is a highly specialized cellular mechanism that not only clears dead cell mass, but also initiates a regulatory immune response to avoid inflammatory tissue damage and autoimmunity [1-2]. Primarily, this function is performed by professional phagocytes such as macrophages. Although in some tissues, nonprofessional phagocytes like endothelial cells can also execute the process [1].

Defects on the clearance of apoptotic cells have shown to affect disease progression, like autoimmune disease and cancer [3]. Macrophage efferocytosis arguably plays the most significant role in apoptotic cell clearance throughout the body. In the liver, the hepatic macrophages are essential in clearing damaged hepatocytes; in the spleen and liver, macrophage efferocytosis is critical for the clearance of dying red blood cells; in the lung, alveolar macrophages are responsible for the clearance of infected pulmonary cells and activated neutrophils through efferocytosis to maintain lung homeostasis; in the blood vessel, macrophage efferocytosis is important to prevent atherosclerosis [4]. Altogether, defects in macrophage efferocytosis can lead to severe health complications.

Mechanistically, efferocytosis is a specialized form of phagocytosis in which the efferocytic cells recognize and engulf only the apoptotic cells for clearance. There are three main stages of efferocytosis which includes, the “find-me”, “eat-me”, and “post-engulfment” stages [5]. The “find-me” and “eat-me” stages are dependent on the direct interactions between the apoptotic and macrophage cells. When cells undergo apoptosis, the apoptotic cells will release chemoattractants, such as nucleotides UTP and ATP, to mobilize macrophages to the site of cell-death [3,5]. The inner membrane leaflet of the apoptotic cells will also flip outwards towards the microenvironment, exposing the inner membrane leaflet phospholipids, like phosphatidylserine (PS). The exposed PS will then promote cell-to-cell interactions through direct and indirect interactions between PS and receptors present on the macrophage. Several receptors have been shown to be involved in the “eat-me” stage of efferocytosis [5].

Of those receptors, the TAM family receptor tyrosine kinases (TYRO3, AXL, MERTK) and their ligands, growth-arrest specific 6 (GAS6) and Protein S (PROS) are the essential regulators, not only in engaging the “eat-me” process, but also in relaying downstream signaling to initiate anti-inflammatory responses to circumvent potential tissue damage signal released by apoptotic cells [6]. Consequently, dysregulated TAM family receptors have been associated with autoimmune diseases and cancer [7-8], making the TAM family receptors promising therapeutic targets to modulate efferocytosis [9]. To engage efferocytosis, the TAM family receptors are activated by their associated ligands GAS6 and PROS, which act as a bridging molecule between PS and the TAM family receptors [6]. Structurally GAS6 and PROS have a Gla domain at the N-terminus, which binds to PS in a calcium-dependent manner, and two LG domains in the C-terminus, which are responsible for interacting with the Ig1-Ig2 domains of the TAM family receptors for activation [6].

Despite current knowledge of TAM family receptors and their relation to efferocytosis, many questions still need to be answered to fully understand how TAM family receptors regulate macrophage efferocytosis in tissues under different physiological or pathology conditions. Addressing these questions will require experiments involving isolated macrophage in culture. Although primary macrophages have been useful for efferocytosis studies [10-16], deriving primary macrophages can be tedious, labor intensive and have high batch-to-batch variation, which all can lead to increase complexity of setting up consistent experiments. A macrophage cell line system would therefore be a useful tool for efferocytosis experiments *in vitro* for the advantages of consistent doubling times, being immortalized and easy to passage, and a uniform genetic background minimizing variability [17].

Currently, there are a few human macrophage cell lines available. The THP-1 cell line has been shown to exhibit observable phagocytic activities and has been widely used to represent human monocytes/macrophages and in phagocytosis studies. Although THP-1 cells have also been used for apoptotic cell engulfment experiments to mimic efferocytosis [11,18,19], careful characterization has not been carried out to validate whether THP-1 cells exhibit specific efferocytosis activities that are distinguishable from phagocytosis. In addition, despite similarities shared by both THP-1 cells and primary macrophages [17], it has been previously noted that THP-1 derived macrophages do not entirely emulate macrophage response to some activating stimuli [20]. Whether those differences would affect efferocytosis experiments carried out with THP-1 cells or not is therefore important to researchers attempting to utilize the THP-1 cell line. In consideration of this, we examined the differential expression of the TAM family receptors and the ligands GAS6 and PROS during THP-1 differentiation states. We also engineered and validated a synthetic, controllable silica bead mimetic assay for efferocytosis activity characterization. Altogether we validated THP-1 cells a reliable human macrophage cell line suitable for efferocytosis experiments.

## Results

### THP-1 differentiation and polarization

THP-1 is a monocytic leukemia cell line (Mo) that is widely used to model human monocyte and macrophage states in culture. The advantage of using THP-1 cells is their plasticity to be induced into differentiated, uncommitted (MΦ) and polarized (M1, M2) macrophages upon treatments [17]. In brief, MΦ differentiation can be achieved by stimulating THP-1 monocytes with phorbol-12-myristate-13-acetate (PMA) for an incubation period (PMA-Phase), followed by a period in culture media without PMA (Recovery-Phase; Fig 1A). The resulting MΦ cells can be further polarized into M1 and M2 macrophages by stimulating the MΦ cells with IFNγ/LPS for 24 h and IL4/IL13 for 48 h respectively (Fig 1A) [22-25].

**Figure 1.**
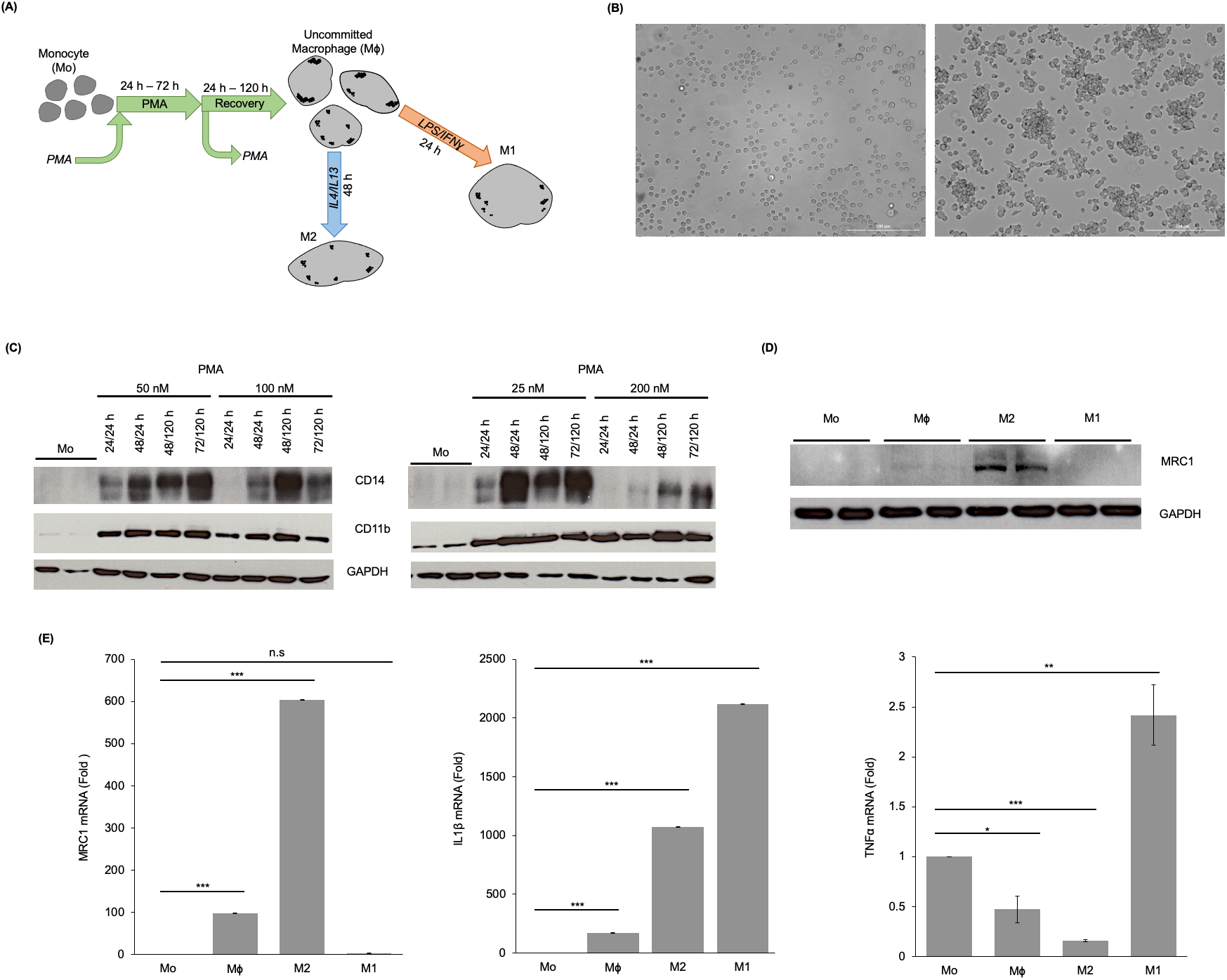
Establishing the phenotypic characteristics of differentiated and polarized THP-1 derived macrophages. **(A)** Representative diagram for differentiating and polarizing THP-1 monocytes into macrophages. THP-1 monocytes were first differentiated into uncommitted macrophages (MΦ) using phorbol 12-myristate 13-acetate (PMA) and were further polarized into either M1 or M2 macrophages by the addition of IFNγ/LPS or IL4/IL13, respectively. **(B)** Brightfield images displaying morphological characteristics of THP-1 monocytes (Mo; Left) and undifferentiated (MΦ; Right). Scale bar, 100 μm. **(C)** Western blot analyses of macrophage differentiation markers. Varied PMA concentrations and incubation periods were tested to establish an optimal differentiation protocol defined by MΦ macrophage CD markers. **(D)** Western blot analyses of M2 macrophage MRC1. Cells were treated with 50 nM PMA for 48 h followed by a 24 h recovery period, then polarized with IFNγ/LPS (M1) or IL4/IL13 (M2). **(E)** RT-qPCR analysis of MRC1, TNFα, and IL1β mRNA. mRNA results are representative of technical triplicates. Cells were treated the same way described in (D). Results were normalized using GAPDH and presented as the mean relative to the untreated monocytes. Error bars are representative of the standard error of the mean using ΔΔCt values. * P<0.05, **P<0.01, ***P<0.001 (One-way ANOVA with Tukey HSD). All western blot data displayed are representative of three independent experiments.

Due to the wide range of reported PMA/Recovery conditions used for THP-1 differentiation [17,26-28], conditions for consistent THP-1 characterization were optimized by testing several combinations of PMA/Recovery conditions (Fig 1C). Morphologically, for all conditions tested, the treated THP-1 cells exhibited known characteristics [17,29,30] of MΦ cells: adherent and increased cell size with visible internal vesicles (Fig 1B). We further checked known CD markers of MΦ macrophages [26,28,30-33] and confirmed the increased expression of CD14 and CD11b upon various PMA-induction conditions (Fig 1C). It is worth of mentioning that the change in CD14 expression was most notable when following incubation periods longer than 48/24 h (PMA/Recovery) with a PMA concentration below 100 nM (Fig 1C). This finding is consistent with data outlined by Mae et al. [27], supporting the use of reduced concentrations of PMA to stimulate THP-1 cells in macrophage differentiation and polarization studies. Overall, although we found that various differentiation conditions can yield MΦ macrophages, for rapid generation of MΦ macrophages with limited unwanted cellular complications by the PMA treatment, we used the protocol of 50 nM PMA, 48 h incubation period (PMA-phase) followed by a 24 h Recovery-phase as the ideal condition for macrophage differentiation.

To further derive M1 and M2 polarized macrophages, we further stimulated the cells with IFNγ/LPS for 24 h and IL4/IL13 for 48 h respectively. Under these treatment conditions, increased expression of the standard M2 marker CD206/mannose receptor (MRC1) was confirmed in the IL4/IL13 treated cells (Fig 1D, 1E) whereas the increased expression of known M1 markers IL1β and TNFα were confirmed in the IFNγ/LPS treated cells. Notably, there was a general increase in IL1β expression for both polarized THP-1 macrophages although a much higher expression was seen in M1-polarized macrophages (Fig 1E).

### Differential expression of TAM family receptors and GAS6 during THP-1 MΦ differentiation

After confirming the differentiation and polarization conditions of the THP-1 cell line, we next investigated the expression patterns of the TAM family receptors during the course of THP-1 macrophage differentiation and polarization. Although some differential expression of MERTK and AXL in bone-marrow derived macrophages (BMDMs) has been reported prior [16], comprehensive characterization of AXL, TRYO3 and MERTK during the phases of macrophage differentiation and polarization has not been thoroughly described, especially within the THP-1 cell line. Since the TAM family receptors have been implicated to being key mediators of efferocytosis, we aimed to explore their expression patterns during the PMA-induced differentiation and polarization phases.

After PMA treatment, AXL and TYRO3 expression were shown to be downregulated, whereas MERTK expression was upregulated (Fig 2A). This differential expression pattern, specifically AXL and MERTK, was consistent with previous studies within BMDMs [16], where AXL expression correlated with inflammatory stimuli and MERTK correlated with tolerogenic stimuli. The upregulation of MERTK from the monocytic THP-1 cells to differentiated macrophage suggests that MERTK is possibly the predominant TAM family receptor in mediating macrophage efferocytosis within THP-1 macrophages.

**Figure 2.**
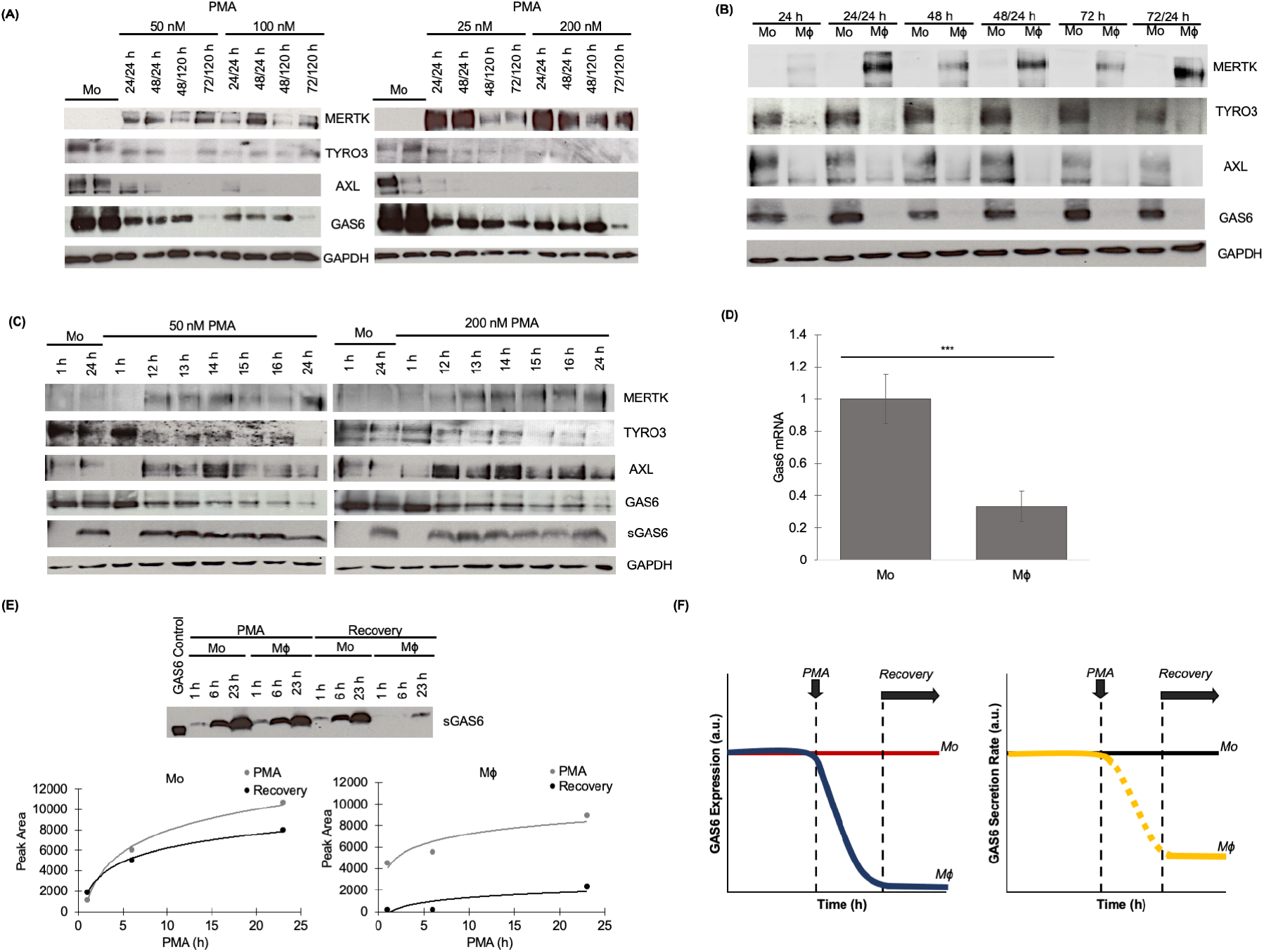
PMA-induced differentiation elicited expression changes of the TAM receptors and the ligand GAS6. **(A)** Western blot analyses of the TAM family receptors and ligands in differentiated THP-1 (MΦ). **(B)** TAM family receptors and GAS6 protein levels were analyzed from cells treated with 200 nM PMA for 24, 48, 72 h alone or followed by a certain recovery period (for example, 24/24 = 24 h PMA followed by 24 h recovery). **(C)** A time-course analysis of the expression of the TAM family receptors, cytosolic GAS6, and secreted GAS6 (sGAS6) during the PMA phase of the THP-1 differentiation protocol. Cells were treated with 50 nM or 200 nM PMA. Total cell lysates and culture medium were collected 1, 12-16, and 24 h after the introduction of PMA. **(D)** RT-qPCR analysis GAS6 mRNA level from cells treated with the 50 nM PMA 48/24 h differentiation condition. Results are representative of technical triplicates. Results were normalized using GAPDH and presented as the mean and relative to the untreated monocytes. Error bars are representative of the standard error of the mean using ΔΔCt values. *** P < 0.001 (Two-Sample T-test). **(E)** Time-course analysis of GAS6 secretion from THP-1 cells untreated (Mo) or treated with 200 nM PMA. The cell culture media were collected at the specified time points. To observe the secretion of Gas6 during only the Recovery phase, the cell culture media were replaced at the beginning of the Recovery phase with fresh media to remove all of the GAS6 accumulated during the PMA phase. (Left) Mo cells. (Right) Mϕ cells. **(F)** Hypothetical graph of GAS6 expression and secretion level change throughout Mo to Mϕ differentiation. All western blot data displayed are representative of three independent experiments.

In addition to the TAM family receptors, we also examined the expression pattern of the ligands GAS6 and PROS during THP-1 differentiation and polarization. The monocytic THP-1 cells expressed abundant GAS6 protein (Fig. 2A). However, upon differentiation, the GAS6 protein level was found to decrease significantly (Fig 2A). In contrast, PROS showed minimal expression in both the monocytic and macrophage THP-1 cells (S2 Figure). Although, PROS is shown to express higher in Mo, upon differentiation, the expression is shown to downregulate to undetectable levels (S2 Figure). Due to the minimal protein level of PROS, the expression pattern was not further explored. Overall, the expression pattern of the TAM family receptors and the ligand GAS6 was consistent across various PMA-mediated differentiation conditions tested, indicating that the general expression pattern of TAM family receptors and GAS6 seen is associated with macrophage differentiation.

To better understand the differential expression pattern, we explored the two phases (PMA vs. Recovery) of THP-1 differentiation (Fig 2B). Overall, the expression changes of AXL, TYRO3, MERTK and GAS6 were observable during the first 24 hours after PMA stimulation and appeared independent of the length or duration of the Recovery-phase (Fig 2B). To ask whether we could determine the specific timepoint of expression change, we carried out a time-course experiment during the first 24 hours of PMA induction. In general, the change in expression for each TAM family receptors occurred between 1-16 h, except for AXL (Fig 2C). A distinct time-point in which AXL expression decreased was not able to be drawn due to the lack of a consistent pattern. Nevertheless, PMA concentration within the 50 nM – 200 nM range did not affect the expression pattern significantly (Fig 2C). Noticeably, only the cellular GAS6 protein level (total cell lysate fraction) was found to significantly decrease whereas the secreted GAS6 (sGAS6) protein level (the culture medium fraction) was unchanged (Fig 2C). This indicates that PMA treatment caused a downregulation of GAS6 at the total cellular protein level but not in the cumulative level of secreted GAS6 (sGAS6). Further examination of the GAS6 mRNA level (Fig 2D) also confirmed that the decrease in GAS6 seen in the total cell lysate fraction was indeed due to a reduction in GAS6 gene expression and not simply due to post-transcriptional regulation of the protein. Altogether, it is possible that GAS6 autocrine signaling is cell differentiation-associated and thereby is regulated in a cell autonomous manner. To further address the GAS6 secretion level change through THP-1 differentiation, we collected the culture supernatant at different time points during the PMA and Recovery phases, and analyzed the protein level of secreted GAS6 (Fig 2E). It was clear to us that GAS6 secretion was already dropped to a very low level at the beginning of the Recovery phase during differentiation (Fig 2E). Given that GAS6 downregulation already occurred during the PMA phase, this decreases in secretion most likely had happened prior to the entry of the Recovery phase (Fig 2E, 2F). In Mo cells, GAS6 is highly expressed, which permits the accumulation of the protein within the cells as well as secreted protein in the supernatant. Once differentiated into MΦ, the downregulation of GAS6 expression caused the remaining cellular GAS6 to be eventually depleted after secretion. We therefore were able to establish a relationship between total and secreted GAS6 during THP-1 differentiation.

### Differential expression of TAM family receptors and GAS6 expression in M1 and M2 polarized THP-1 cells

Most effector macrophages in tissues are often polarized upon infection, inflammation or tissue damage. To characterize the physiological relevance of using the THP-1 cell model system for functional efferocytosis applications, we further examined the expression of the TAM family receptors within the standard M1 and M2 phenotypes. Following the scheme described in Fig. 1A, M1 or M2 polarized macrophages were generated by stimulating the THP-1 MΦ with IFNγ/LPS (for M1 polarization) and IL4/IL13 (for M2 polarization). Overall, the decrease in TYRO3 seen in MΦ cells was seen in both in M1 and M2 polarization, implying that TYRO3 signaling does not have a significant role in M1 and M2 macrophage efferocytotic activity (Fig 3A). In contrast to TYRO3, both

**Figure 3.**
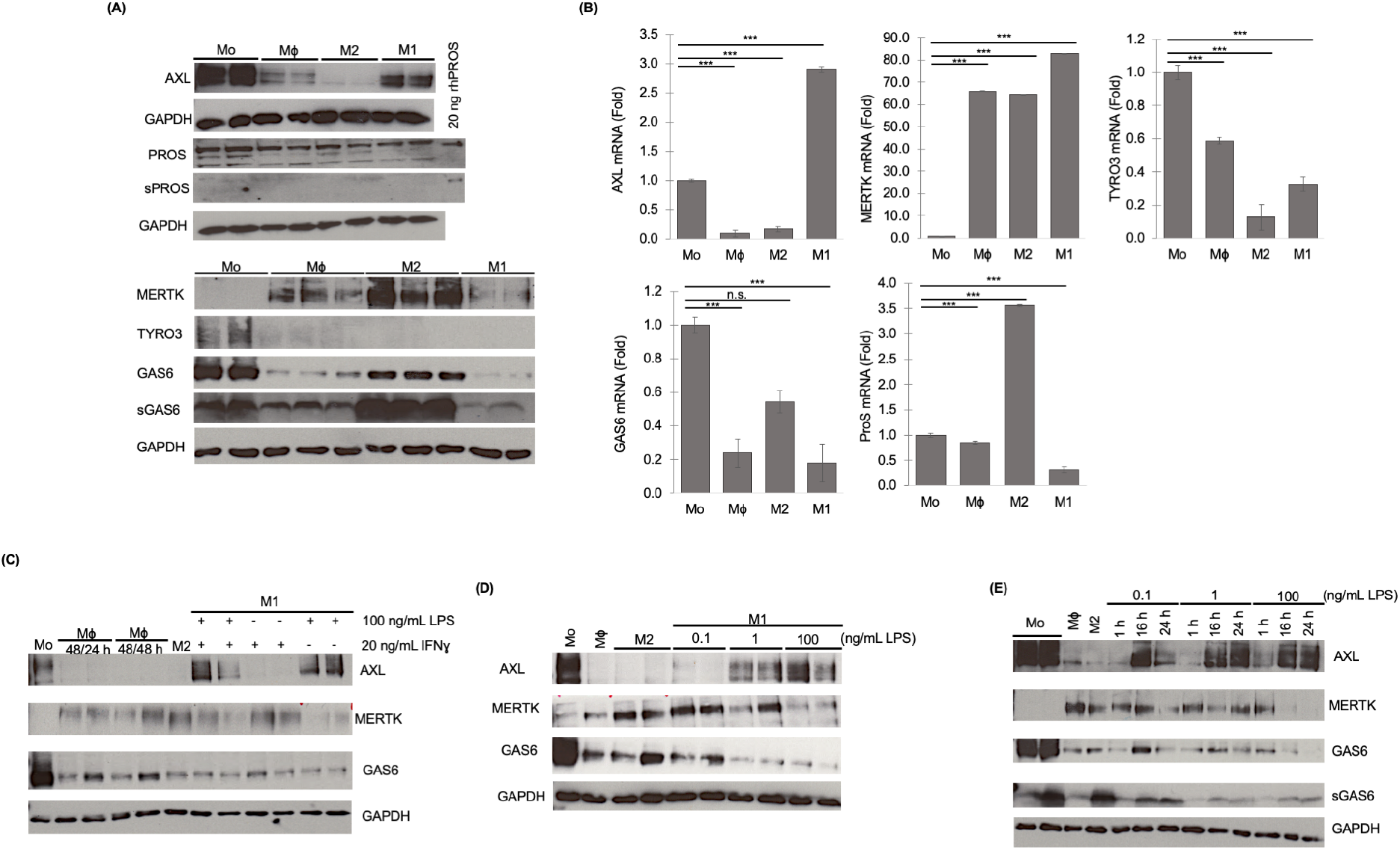
The effect of polarizing THP-1 derived macrophages into M1/M2 phenotype on TAM receptor and associated protein expression. Mϕ THP-1 cells were derived following the 50 nM PMA 48/24 h differentiation protocol. Cells were then further polarized into either M1 or M2 macrophages. Total cell lysates and cell culture supernatants were collected at the end of each treatment period and analyzed by western blots. **(A)** TAM family receptors, GAS6/PROS and secreted GAS6/PROS (sGAS6/sPROS) expression in Mo, Mϕ, M1, and M2 THP-1 cells. **(B)** RT-qPCR analysis TAM receptors and TAM ligands mRNA level from cells treated with the 50 nM PMA 48/24 h differentiation condition. Results are representative of technical triplicates. Results were normalized using GAPDH and presented as the mean and relative to the untreated monocytes. Error bars are representative of the standard error of the mean using ΔΔCt values. n.s. P > 0.05, *** P < 0.001 (One-way ANOVA with Tukey HSD). **(C)** TAM family receptors, GAS6, and secreted GAS6 (sGAS6) expression in THP-1 treated with either LPS, IFNγ, or in combination. **(D)** Cells were treated with varying LPS concentrations in combination with 20 ng/mL IFNγ for 24 h. **(E)** Time course (1, 16, and 24 h) analysis of cells treated with 0.1, 1, or 100 ng/mL LPS combined with 20 ng/mL IFNγ. All western blot data displayed represent **(A-B)** three or **(C-D)** two independent experiments.

MERTK and AXL showed differential expression patterns among the MΦ, M1 and M2 cell types. AXL expression was reduced to the greatest extent in M2 cells, with significant reduction seen in MΦ cells, as well as slight reduction in M1 cells (Fig 3A). There was a notable increase in MERTK in all three cell states, although with M2 showing the greatest increase in protein level (Fig 3A). In addition, we saw a significant rebound of GAS6 and PROS in M2 cells (Fig 3A). The expression patterns were further validated at the mRNA level of each protein. The expression patterns of AXL, TYRO3, PROS and GAS6 all showed consistent findings with the Western blot data (Fig 3B). However, MERTK patterns were not consistent between the mRNA and the observed protein level (Fig 3B). At the mRNA level, there is a general increase in MERTK expression, which is consistent with our western blot results. However, when exploring between the macrophage cell-types, the patterns between mRNA expression and protein levels do not match. M1 macrophages showed the highest expression of MERTK, however, at the protein level MERTK expression is the lowest amongst the three-macrophage cell-types (Fig 3A & B). It is likely that this discrepancy between the mRNA and protein levels is due to regulation between mRNA levels and protein transcription. Further exploration is required to elucidate the reasoning behind this variation in MERTK mRNA and the protein levels.

Interestingly, among the MΦ, M1 and M2 cell types we found that AXL expression correlated well with the M1 cell-type, suggesting that AXL can potentially be a useful marker for M1 macrophages. To examine the effect of the M1 polarization stimuli on AXL expression, we used varied polarization conditions to explore whether the expression pattern can be affected by any of the M1 polarization stimulants. By removing either LPS or IFNγ, we identified that LPS, and not IFNγ, was the determinant factor responsible for the differential expression of AXL and MERTK (Fig 3C). The effect of other IFN proteins on TAM receptor expression were also further explored, as Type I IFNs are known regulators of the TAM kinase receptors [34,35]. As expected, increased concentration of IFNβ and IFNα also increased AXL expression (S3 Figure). TYRO3 expression was also shown to be affected by IFNβ, in which TYRO3 expression decreased as the concentration of IFNβ increased (S3 Figure). MERTK expression was not affected by either IFNβ or IFNα.

We next performed M1 polarization with varied doses of LPS while keeping the IFNγ concentration fixed at 20 ng/mL. THP-1 macrophages were treated with either a low (0.1 ng/mL), intermediate (1 ng/mL), or high dose (100 ng/mL) of LPS following the standard M1 polarization procedure. AXL expression was shown to increase in a dose-dependent manner, with doses greater than or equal to 1 ng/mL (Fig 3C). Accompanied with the elevated AXL level, a decrease in both MERTK and GAS6 expression was also observed (Fig 3C). This observation is in agreement with previous work on murine BMDMs, which showed that LPS alone induced the increase in AXL and decrease in MERTK expression [16]. This similarity indicates that the THP-1 cell line can recapitulate the features of primary BMDMs and further suggests the usefulness of the THP-1 cell line as a model system for macrophage studies. By setting up a time-course experiment, we further confirmed that AXL upregulation occurred within the first 16 hours of M1 polarization (Fig 3E). We also observed a strong suppression of MERTK and GAS6 expression after exposure to concentrations greater than or equal to 1 ng/mL of LPS (Fig 3E). These findings could potentially be exploited to use AXL as a M1 polarization indicator.

Altogether, we have profiled the expression patterns of the TAM family receptors in M1 and M2 THP-1 macrophages. Based on the upregulated expression levels of AXL and MERTK, we speculate that both receptors play major roles in dictating polarization-related efferocytic function. The differential expression of TAM family receptors as well as associated ligands in M1 and M2 would also suggest that each receptor, notably AXL and MERTK, have varied roles. Since the M1 macrophage is mostly induced by infection, AXL expression could be correlated with efferocytic activities associated with inflammatory response. In contrast, MERTK as the predominant TAM family receptor in M2 may be involved in efferocytosis associated with immune tolerance and tissue repair. The low expression of TYRO3 in all three macrophage cell types may indicate that TYRO3 does not have a significant role in THP-1 macrophage function. Furthermore, the association of GAS6 and MERTK expression pattern in M2 phenotype indicates that THP-1 cell exhibits distinct properties of GAS6 autocrine signaling loop among the three cell states (MΦ, M1, M2), which may serve important regulatory functions in cell state-specific efferocytosis.

### Evaluating efferocytosis activity in THP-1 derived macrophages

To evaluate the utility of THP-1 derived macrophages for efferocytosis studies, we next set up a quantitative efferocytosis assay. First, to derive apoptotic cell fragments for inducing efferocytosis, we generated an inducible apoptosis HEK293 cell line expressing a chemical-activated caspase 9 (iCasp9) (Fig 4A). This system is composed of a modified caspase-9 protein, in which the CARD domain is replaced by a FKBP12-F36V domain, allowing inducible caspase-9 dimerization upon introduction of the synthetic FKBP12-F36V ligand AP20187 [36,37] to initiate apoptosis. To confirm the successful induction of apoptosis in this system, we used cleaved-PARP and phosphatidylserine (PS) as markers. An increase in cleaved-PARP was evident as early as 2h after induction with AP20187 (Fig 4B). Additionally, using fluorescently labeled Annexin V as a detection agent [38], the cell surface exposure of PS was evident after a 4h induction detected (Fig 4C). These findings altogether demonstrated the successful apoptotic cell generation using the iCasp9 HEK293 cell line.

**Figure 4.**
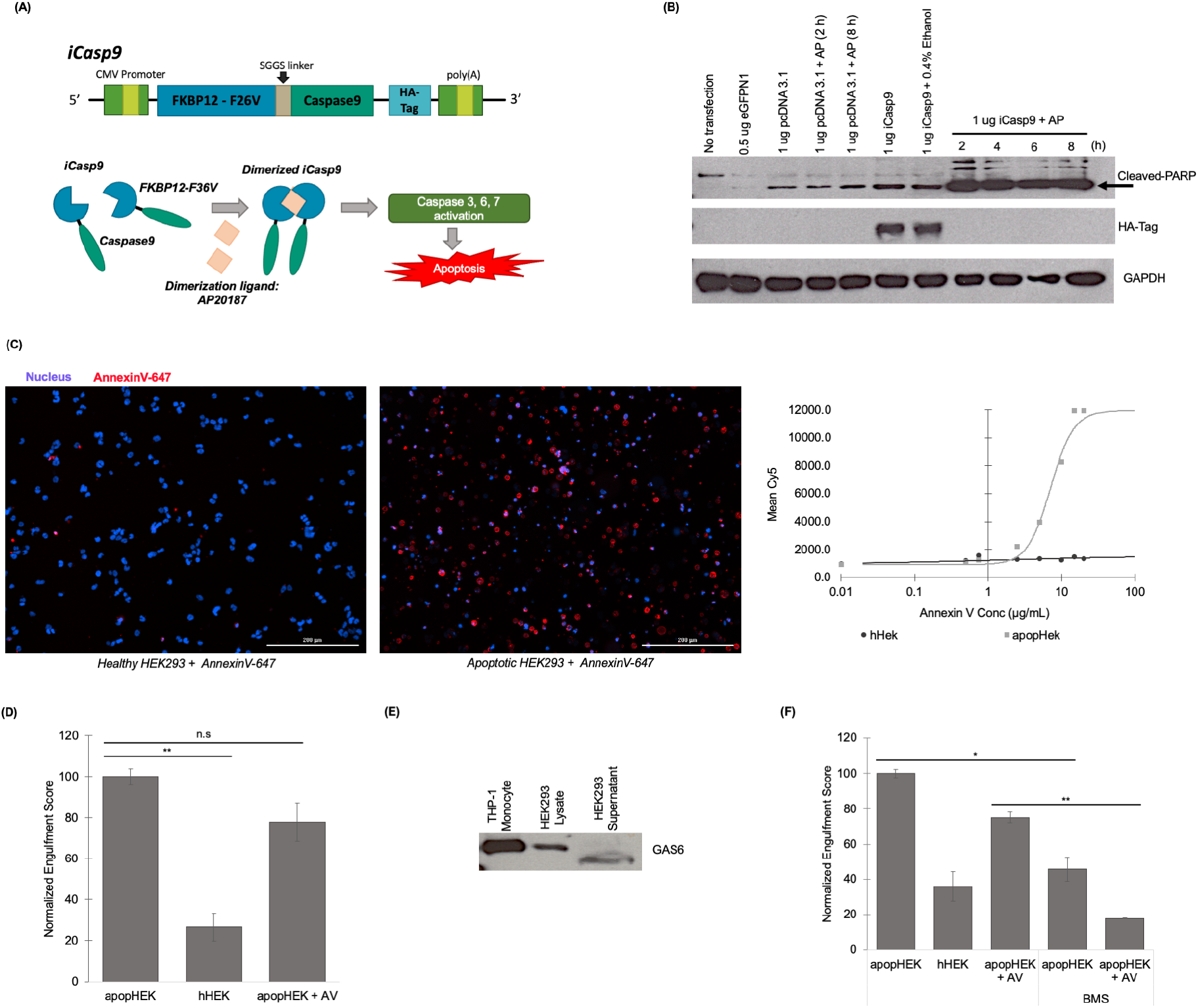
TAM-mediated efferocytosis potential of THP-1 derived macrophages. **(A)** Representative diagram of the iCasp9 inducible system to induce apoptosis. iCasp9 HEK293 cells were treated with 1 μM AP20187 for 4 h to induce apoptosis. **(B)** Total cell lysate samples were collected and analyzed using western blot analysis. **(C)** Apoptotic (apopHEK) and healthy (hHEK) iCasp9 HEK293 cells were collected and incubated with 15 μg/mL Annexin V-647 to visualize and quantify the cell surface PS. Cells were also stained with 1 μg/mL Hoechst 33342 to label the nucleus. PS level was quantified by measuring the Annexin V-647 fluorescent intensity of each image. Scale bar, 200 μM. **(D)** THP-1 cells were differentiated into Mϕ macrophages and were subsequently exposed to either apopHEK, hHEK, or Annexin V-blocked apopHEK for a 2 h co-incubation. **(E)** Total cell lysate and cuture medium fractions were collected and analyzed by western blot analysis. **(F)** THP-1 Mϕ macrophages pre-treated with 5 μg/mL BMS-777607 prior to a 2 h co-incubation with either apopHEK, hHEK, or Annexin V-blocked apopHEK. All efferocytosis data was imaged using fluorescent and brightfield microscopy and further analyzed using ImageJ software (S1 Protocol). Each experimental setup was repeated as **(E-F)** two or **(B-D)** three independent experiments. Error bars are representative of the standard error of the mean. (D & F) Welch’s ANOVA with Games-Howell Post Hoc Test. n.s. P > 0.05, * P<0.05, ** P < 0.01, *** P < 0.001.

Using quantitative image analysis (S1 Methods) to quantitate internalized cells, we found that MΦ cells fed with apoptotic HEK293 cells (apopHEK) showed a significant level of engulfment whereas MΦ cells fed with healthy HEK293 cells (hHEK) showed minimal engulfment (Fig 4D). The significant differences between apopHEK and hHEK engulfment indicated that the apoptotic cells indeed triggered measurable efferocytosis in THP-1 MΦ cells.

In general, there are two routes that apoptotic cells can engage macrophage efferocytosis: one is through the direct interaction of PS and the PS-recognizing receptors (TIM1/4, BAI or Stabilin) [5,39]; the other is through the TAM family receptors, where the ligands GAS6/PROS bridge PS and the TAM family receptors. For both scenarios, the presence of PS is required, and in principle, by masking PS with PS-binding molecules, efferocytosis can be blocked, permitting only phagocytosis. To elucidate that the efferocytosis seen was mediated by TAM family receptors, we used Annexin V as a PS-blocking molecule to attenuate the interaction between PS and cell surface PS receptors (Fig 4D).

Adding Annexin V to apopHEK cell fragments, we observed a slight decrease in apoptotic body engulfment (Fig 4D; apopHEK + AV), indicating that the efferocytosis activity was partly contributed by PS receptors on THP-1 derived macrophages. Nevertheless, the fact that Annexin V did not fully abolish apopHEK efferocytosis suggested that there was still a form of efferocytosis activity occurring, most likely mediated by the TAM family receptors. The expression and secretion of GAS6 (Fig 4E) also supported the notion that apopHEK cell fragments could contain GAS6 to activate TAM family receptor-mediated efferocytosis. To test this a pan-TAM kinase receptor inhibitor, BMS-777607, was used. MΦ cells treated with BMS-777607 showed a significantly decreased efferocytosis, demonstrating that the TAM family receptors are as the major efferocytosis effector for apopHEK cell fragments (Fig 4F). Moreover, addition of Annexin V also further decreased the efferocytosis level, in agreement with the previous finding shown in Fig 4 E. Overall, we found that MΦ cells showed a significant decrease in engulfment upon blocking general and TAM-mediated efferocytosis activity (Fig 4F), providing evidence that the THP-1 derived macrophages exhibit notable efferocytosis activities, thus, making THP-1 a suitable cell line for efferocytosis studies.

### Synthetic apoptotic body mimetics to assay efferocytosis

Although feeding macrophages with apoptotic cells is the common practice for measuring efferocytosis [10,12,16,19,40,41], there are some drawbacks in using apoptotic cells to assay efferocytosis experimentally. The major concern was that the protein contents of the apoptotic cells are less well-defined, and the secretory profiles from some cell lines can contain undefined levels of GAS6 and PROS, as was seen with the HEK293 cells, making the efferocytosis interpretation less consistent and the assay more convoluted. We therefore sought to explore the use of a synthetic assay system for quantitating efferocytosis. We thus strategized an approach of using synthetic silica beads to mimic apoptotic cells, with defined protein and lipid components on the beads. Phosphatidylserine (PS) is a surface landmark that is featured on the outer leaflet of apoptotic cells, whereas phosphatidylcholine (PC) represents the external membrane leaflet of healthy cells. We therefore generated fluorescently labeled PS-conjugated beads to mimic the apoptotic cells or PC-conjugated beads to mimic healthy cells as a comparison. The chemical-defined fluorescent beads thus provided us the full controllability to dissect the efferocytic properties of THP-1 cells.

To examine whether THP-1 cells could respond to the apoptotic body mimetics, we first fed the PS-beads to differentiated THP-1 MΦ macrophages. At the completion of the 2 h incubation, the cells were extensively washed followed by confocal microscopy imaging. Using quantitative image analysis (S1 Methods) to quantitate internalized beads, we were able to visualize a significant increase in bead engulfment over the 2 h incubation (Fig 5A). Since efferocytosis can be considered as a specialized form of phagocytosis responsible for the clearance apoptotic cells, macroscopically these two processes (i.e., phagocytosis vs. efferocytosis) do share several common features in cellular engulfment. To distinguish if the engulfment seen was efferocytosis or phagocytosis, in addition to making PS-beads, we also introduced PC-beads to THP-1 MΦ macrophages. To our expectation, there was an observable engulfment of PC-beads in THP-1 macrophages (Fig 5B). However, as compared to the level of PS-beads engulfment, the level PC-beads engulfment was significantly lower. We therefore concluded that the level of PC-beads engulfed can be set as the basal level of THP-1 phagocytosis, whereas the PS-beads engulfment can be marked as efferocytosis or in combination with some phagocytic activity. Similar to what was done for the apopHEK cells, we blocked the PS-beads with Annexin V, showing saturation levels around 0.736 μg/mL of Annexin V (S4 Figure) and found that cells fed with Annexin V-blocked PS-beads (PS-AV) showed a significant decrease in engulfment similar to that of cells fed the PC-beads (Fig 5B). This result is also consistent with previous findings which found that PS is key in the recognition of the apoptotic cell to induce efferocytosis [42]. Moreover, using chemical-defined beads and controllable ligands, we were able to distinguish efferocytosis from phagocytosis with complete control of the system.

**Figure 5.**
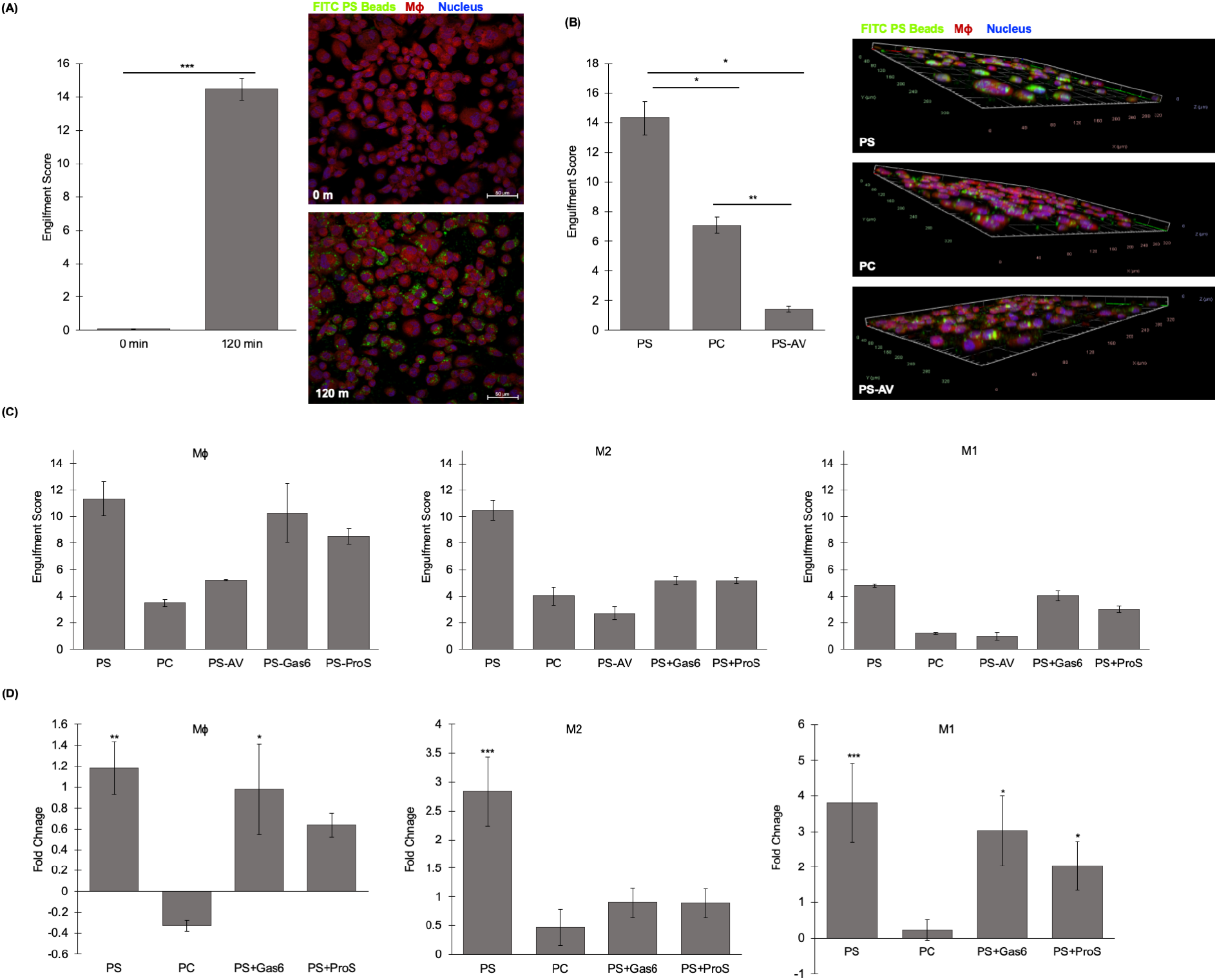
Efferocytosis activities of THP-1 derived macrophages. **(A)** Confocal microscopy of Mϕ THP-1 cells fed with FITC labeled PS-beads for either less than 1 min (top) or 120 min (bottom). CP 670 (red) and Hoechst 33342 (blue) were used to stain the whole cell and nucleus, respectively. Scale bar, 50 μM. Quantitative engulfment score from ImageJ analysis (S1 Protocol) of images. **(B)** Mϕ cells fed with PS- or PC-beads, preloaded with and without Annexin V (PS-AV), for 120 min. Z-stack images were used to confirm engulfment. **(C)** Mϕ, M1, and M2 THP-1 cells fed with PS-beads, PC-beads, or PS-beads decorated with saturating amount of Annexin V, Gas6 (PS-Gas6), ProS (PS-ProS) for 120 min. **(D)** Fold change of engulfment score. The fold change was calculated relative to the engulfment score of PS-AV for each cell type. Data is representative of technical triplicates. Each experimental setup was repeated as three independent experiments. Error bars are representative of the standard error of the mean. (B) Welch’s T-test. (C & E) Welch’s ANOVA with Games-Howell Post Hoc Test. * P<0.05, **P<0.01, ***P<0.001. All conditions were imaged by confocal microscopy and further analyzed using ImageJ software.

To assess TAM-mediated efferocytosis, we introduced GAS6 and PROS ligands to the PS-beads to engage TAM family receptors for efferocytosis. To ensure the engulfment only involves TAM family receptors and not PS receptors, we used high amounts of GAS6 and PROS to fully occupy the bead surface PS, based on the equal molar ratio of the Annexin V saturation curve (S4 Figure). Structurally, the PS binding domain of GAS6/ PROS is analogous to that of the Annexin V. Therefore, similar to PS-AV beads, when the PS-beads were preloaded with saturating amounts of GAS6/PROS, all PS would be masked and excluded from the direct interactions with the PS-receptors. However, unlike Annexin V, the receptor binding domains of GAS6 and PROS can engage the beads to the TAM family receptors, thereby permitting only TAM-mediated efferocytosis. In this set up, THP-1 macrophage types (MΦ, M1 and M2) did show varying degrees of GAS6/PROS-PS-beads engulfment compared to the PS-AV control. (Fig 5C). We therefore concluded that by controlling the ligands present on the PS-beads we are able to switch THP-1 efferocytic activities from PS receptor-mediated to TAM family receptor-mediated.

Among the three macrophage types, M1 macrophage generally exhibited a significantly lower efferocytic activity (Fig 5C). As noted, when exposed to either the GAS6 or PROS saturated PS-beads, the TAM family receptor mediated-efferocytic activities seemed to be much lower to that of PS receptor mediated in M2 macrophages. Further looking at the fold change of bead engulfment in relation to cells exposed to PS-AV (Fig 5D), it was more obvious to see the receptor usage preference of efferocytosis in different macrophage types. In M1 and MΦ cells, TAM family receptor and PS-receptor mediated efferocytic activities are comparable. However, in M2 cells only the PS receptor mediated efferocytic activity was significant. It is therefore intriguing that M2 macrophages has a higher preference in PS-directed over the TAM family receptor-mediated efferocytic pathway, despite MERTK expression level is high. Our assay set up using chemical-defined beads and controllable ligands thus provided a valuable tool for the identification of this phenomenon.

## Discussion

Efferocytosis plays an essential role in maintaining homeostasis by removing dying cells from tissue. The regulation of this biological process is complicated. Dysregulated efferocytosis have been implicated in multiple diseases, including autoimmune disorders, atherosclerosis and cancer [1,3,39]. In particular, aberrant functions of TAM family receptors in macrophage have been shown closely associated with dysregulated efferocytosis in animal models and in human [6,43]. Understanding the molecular mechanisms by which TAM family receptors regulate efferocytosis is therefore significant and could lead to novel therapies. To provide an *in vitro* culture system for cellular and biochemical characterization of efferocytosis, we characterized the THP-1 macrophage and demonstrated its usefulness in representing primary BMDMs for efferocytosis studies.

In this study we first described the differential expression of the TAM receptors amongst the undifferentiated THP-1 monocytes and the different macrophage. Most notable was the shift in expression of AXL and MERTK during macrophage differentiation and polarization. TYRO3 expression, however, was not abundantly detected in all THP-1 cells. It is possible that TYRO3 has less significance in macrophage and dendritic cells lineages [35]. Interestingly, THP-1 monocytes express high levels of GAS6. However, the expression is significantly downregulated during macrophage differentiation and was further suppressed in M1 macrophages. In contrast, GAS6 expression was maintained in M2 macrophage, although notably lower when compared to monocytes. Given the different expression patterns (S2 Table) found in THP-1 monocyte, MΦ, M1 and M2 macrophages, no clear cell-type associated expression correlation can be drawn among GAS6, AXL and MERTK. However, it was obvious that LPS and type I IFNs can affect the expression of AXL, MERTK and GAS6 in M1 macrophage. Whether THP-1 monocytes or macrophages requires GAS6 signaling in an autocrine-regulated manner is await to be investigated in the future. PROS showed very minimal expression in the THP-1 derived macrophages, suggesting PROS is dispensable for THP-1 differentiation and polarization (Fig 3A). Nevertheless, the effect of PROS as a stimulating factor for macrophages should be further explored to better understand its role in immune response.

Notably, THP-1 macrophages exhibit significant efferocytic activities. More importantly, we confirmed that the THP-1 macrophages have higher efferocytic activity than general phagocytic activity, making it a valid cell line for *in vitro* efferocytosis studies. It is also an important feature of our study to evaluate and to distinguish efferocytosis from general phagocytosis. In our assay system, we found two distinct levels of engulfment activity when the THP-1 cells were exposed to GAS6/PROS-bound beads in comparison to PS-beads (Fig 5D).

Cell surface exposed PS has long been known as a surface landmark on apoptotic cells. However, its involvement in initiating efferocytosis was not understood until the discovery of the PS receptors TIM-1 and TIM-4 and ligands GAS6 and PROS [12, 42, 44-48]. The TIM family proteins are glycoprotein receptors including TIM-1, TIM-2, TIM-3 and TIM-4. TIM-1 and TIM-4. They have been known as high affinity PS-binding receptors that directly interact with PS [12]. In contrast, TIM-3 and PS interaction has been proposed [49], but has not been extensively verified. Specifically, TIM4 has been shown to be involved in the process of efferocytosis and is ubiquitously expressed in macrophages [12,50].

With the identification of TIM-4 as a PS-binding receptor and the TAM family proteins, GAS6/PROS, it became clear that efferocytosis can be executed through at least two distinct routes: TIM receptor mediated and TAM family receptor mediated [44-48]. However, how the two distinct routes coordinate or cooperate during efferocytosis still remains to be further investigated. One notable study has shown that within macrophages, TIM-4 is not able to initiate efferocytosis, rather it aids in TAM-mediated efferocytosis [48].

Using the chemical-defined beads, we were able to observe that at least in M2 THP-1 macrophage, TAM family receptors play little role in the engulfment of efferocytosis. Instead, efferocytosis in M2 THP-1 macrophage seemed to be dominated by the direct PS receptor interactions. It has not yet been determined if this general phenomenon would apply to most M2 macrophages. Nonetheless, given that we have shown that THP-1 cells resemble primary BMDMs, it is likely to be the case. Besides TIM-4, a few receptors (BAI1, CD300f, Stabilin) have also been reported as *bona fide* direct PS-interacting receptors that may contribute to efferocytosis [5,39]. However, the presence of these direct PS receptors and their roles within THP-1 derived macrophages has not been thoroughly explored, which we aim to investigate in future studies.

We have demonstrated that THP-1 derived macrophages can be used as a reliable model system to study efferocytosis. We also illustrated the potential of using such a system for screening efferocytosis modulators. Since dysregulated efferocytosis has been implicated in diseases, a robust functional screen system will be valuable for drug discovery. Overall, this study shows that the process of efferocytosis is complex and that additional studies are required to determine receptor interactions, as well as, to distinguish the effect of each receptor on efferocytosis. The THP-1 based efferocytosis assay system can be a valuable tool for such purposes.

## Materials and Methods

### THP-1 Cell Culture

The THP-1 cell line originating from ATCC (American Type Culture Collection) were obtained from validated Cold Spring Harbor Laboratory cell line collection. Cells were cultured in RPMI 1640 media supplemented with 10% heat inactivated fetal bovine serum, 100 U/mL penicillin, and 100 U/mL streptomycin, at 37°C in 5% CO_2_. For Macrophage differentiation, THP-1 monocytes (Mo) were differentiated into macrophages (MΦ) by stimulating cells with a range of concentrations (25 nM – 200 nM) of phorbol 12-myristate 13-acetate (PMA; InvivoGen) for different incubation periods. The incubation periods consist of a PMA and Recovery phase. The PMA phase consists of stimulating the cells with the addition of PMA in the growth media. During the Recovery phase, the PMA-supplemented growth media is removed and replaced with media without PMA to allow the cells to reach a resting state. A differentiation protocol was established for all polarization experiments, which comprises of stimulating the cells with 50 nM PMA for 48 h followed by 24 h. M1 polarized cells were obtained by stimulating MΦ cells with 20 ng/mL of IFNγ (Stemcell Technologies, Cat# 78020) and 100 ng/mL of LPS for 24 h. M2 polarization was achieved by stimulating MΦ cells with 20 ng/mL of interleukin 4 (IL4; Stemcell Technologies, Cat#78045) and interleukin 13 (IL13; Stemcell Technologies, Cat# 78029).

### Western blot analysis

At the conclusion of the differentiation and polarization treatments, whole cell lysates were prepared using cell lysis buffer (10 mM Tris, pH 7.5, 100 mM NaCl, 0.5% sodium deoxycholate, 10% glycerol, 1% Triton X-100 with Pierce protease and phosphatase inhibitor [Thermo Scientific]). Cell culture supernatant samples were also collected. Both lysate and supernatant samples were centrifuged at 16,000 xg at 4 ºC for 10 min. The pellet was discarded and the remaining supernatant was used as whole lysate (lysate) or supernatant samples. For lysates, protein concentrations were measured using a Bradford kit (VWR). The lysate and supernatant samples were mixed with 1x Laemmli buffer with 1.5% 2-mercaptoethanol and were denatured at 95 ºC for 5 min. The samples were separated by a 10% SDS-PAGE and transferred to a nitrocellulose membrane (Amersham Protran 0.45 μm NC; GE Healthcare Life Science). The membranes were blocked in 1x TBS-0.1% Tween-20 (TBS-T) with 5% non-fat dry milk for 30 min at room temperature (RT). Primary antibodies were added and incubated overnight at 4ºC. Before and after adding the secondary antibody, the membranes were washed three times with 1x TBS-T for 10 min each. Membranes were incubated at RT with the secondary antibody for 1 h. The membranes were developed using the x-ray film. Refer to S1 Table for a full list of primary and secondary antibodies.

### Quantitative PCR analysis

Total cellular RNA was isolated for gene expression analysis of IL1β, TNFα, MRC1, MERTK, AXL, TYRO3, GAS6 and PROS. Trizol reagent was used following the Trizol reagent was used following the outlined protocol of the manufacture (Invitrogen Life Technologies). cDNA was synthesized from 2 μg of total RNA using the SuperScript IV Reverse Transcriptase (Invitrogen Life Technologies) protocol. Both Oligo d(T)_20_ and random hexamer primers were used during the synthesis. For the qPCR reactions, the Luna Universal qPCR Master Mix protocol (NEB) was followed.

For amplification of each of the targeted genes, the following primer pairs were used (synthesized by Integrated DNA Technologies): **AXL:** 5’CCGTGGACCTACTCTGGCT (Forward), 5’ CCTTGGCGTTATGGGCTTC (Reverse); **GAS6:** 5’GGTAGCTGAGTTTGACTTCCG (Forward), 5’GACAGCATCCCTGTTGACCTT (Reverse); **GAPDH:** 5’ATGGGGAAGGTGAAGGTCG (Forward) 5’GGGGTCATTGATGGCAACAATA (Reverse); **IL1β:** 5’ATGATGGCTTATTACAGTGGCAA (Forward), 5’GTCGGAGATTCGTAGCTGGA (Reverse); **MERTK:** 5’ACCTCTGTCGAATCAAAGCCC (Forward), 5’CTGCACACTGGTTATGCTGAA (Reverse); **MRC1:** 5’CTACAAGGGATCGGGTTTATGGA (Forward), 5’TTGGCATTGCCTAGTAGCGTA (Reverse); **PROS1:** 5’TCCTGGTTAGGAAGCGTCGT (Forward), 5’CCGTTTCCGGGTCATTTTCAAA (Reverse); **TNFα:** 5’ GAGGCCAAGCCCTGGTATG (Forward), 5’CGGGCCGATTGATCTCAGC (Reverse); **TYRO3:** 5’GAGAGGAACTACGAAGATCGGG (Forward), 5’AGTGCTTGAAGGTGAACAGTG (Reverse).

GAPDH expression was used as a control. Reactions were ran using the QuantStudio 6 Flex Real-Time PCR System (Life Technologies).

### Silica bead preparation

To prepare the bead model used for the efferocytosis assay, 2 μm carboxylic acid functionalized silica spheres (Bangs Laboratory) were functionalized with octadecylamine (Sigma) using DCC coupling chemistry, followed by coating with either 18:1 phosphatidylserine (PS) or 18:1 phosphatidylcholine (PC; Avanti Polar Lipids). <5 molar % FITC (Sigma) conjugated to octadecylamine was added to the lipids to yield fluorescently labeled microparticles. These lipid-coated microparticles were then used for all efferocytosis experiments in this study. To decorate the PS/PC-beads with either Gas6 or ProS, the beads were introduced to either 20 nM recombinant human GAS6 (R&D Systems, Cat# 885-GSB-050), or recombinant human Protein S (R&D Systems, Cat# 9489-PS) with 2.5 mM CaCl_2_ for 1.5 h. To remove unattached protein, the beads were washed twice with 1x PBS, 2.5 mM CaCl_2_ and centrifuged at 3,000 g for 5 min.

### Apoptotic HEK293 production

To generate the iCasp9 HEK293 cell line, HEK293 cells cultured in a 35mm plate was transfected with 1 μg of pcDNA3.1 hyg(+) FKBP-F36V Casp9ΔCARD-HA plasmid. Stable cell lines were established after 200 μg/mL Hygromycin B selection. To induce apoptosis the iCasp9 HEK293 cells were treated with 1 μM AP20187 [Sigma Aldrich] for 4 hrs. Following induction of apoptosis, both adherent and floating cells were collected [apopHEK] and used within the efferocytosis assays. Cell viability was determined using a cell counter and phosphatidylserine (PS) exposure was quantitated by performing an Annexin V binding curve. For the Annexin V binding curve, apopHEK cells were exposed to 0, 0.5, 0.736, 2.5, 5, 10, 15, and 20 μg/mL of Alexa Fluor 647 Annexin V [BioLegend]. Images were attained and analyzed using the BioTek Cytation 5 imaging reader system. Saturation curves of Annexin V were plotted by measuring binding of Annexin V-647 to apopHEK and healthy HEK293 [hHEK] cells.

### Efferocytosis assay

THP-1 cells were cultured in 96-well plates following the 48/24 h differentiation and M1/M2 polarization protocol. At the completion of differentiation and polarization, cells were washed twice with 1x PBS to remove any secreted proteins. The culturing media was reintroduced and either the prepared PS-/PC-beads were added at a ratio of 20:1 (Beads: Cells) or apopHEK cells stained with 5 μM eBioscience CFSE (Invitrogen Thermo Fisher Scientific) were added at a ratio of 5:1 (HEK: THP-1). The cells were incubated with the apoptotic bodies [beads or apoptotic cells; ABs] up to 2 h at 37 ºC. To remove any unattached or unengulfed ABs, the wells were extensively washed with 1x PBS. Cells were fixed using 4% paraformaldehyde for 10 min at RT. THP-1 cells were stained with eBioscience™ Cell Proliferation Dye eFluor™ 670 (Invitrogen Thermo Fisher Scientific) and with Hoechst 33342 (Cell Signaling Technology, #4082).

### Confocal microscopy & image analysis

Cells were imaged using the Zeiss LSM 710 Confocal microscope. Stacked images of the whole cell, nucleus, and the ABs were acquired using the Z-stack feature. The 458 nm (DAPI), 488 nm (FITC), and 633 nm (CP 670) lasers were used to image the cell nucleus, ABs, and cell area, respectively. All confocal images were processed using ImageJ image analysis software [21]. To quantify the overlapping area of the ABs with the cells, the following steps were followed (S1 Method). Images were uploaded using the Bio-Formats importer plug-in. The FITC and CP channel were selected with the appropriate Z-layer (representative of the mid-point of the cell) and duplicate images were produced. The brightness of both duplicate images was increased and the threshold was determined. Using the FITC image, a selection was made (Edit → Selection → Create selection), which highlighted the threshold ABs area. This selection was then transferred to the CP image (shift-e). Under the Analyze tab, measurements were set to count the area and to limit it to the threshold. The CP image alone and with the transferred FITC selection was measured. Using these values, the total overlapping area was determined and normalized against the total cell area in the frame.

### Data analysis

All presented data was done in duplicate or triplicates and were repeated in two or three independent biological experiments. Statistical analyses were performed with IBM SPSS using One-way ANOVA with Tukey HSD (Fig 1), Welch’s T-Test (Fig 2 & Fig 4B), and Welch’s ANOVA with Games-Howell Post Hoc Test (Fig 4C, 4E). Differences with a P value of < 0.05 were considered significant.

## Acknowledgments

We thank Audrey Fahey, Carmelita Bautista, Madison Kallman, Michelle Yang, David Ng for assisting the preparation of reagents. We also thank Fabian Gerth and Craig Podszus for assisting in the preparation of the manuscript. This work was supported in part by Development Funds and the CSHL Shared Resources funded by the Cancer Center Support 5P30CA045508, and the Cold Spring Harbor Laboratory and Northwell Health Affiliation.

