## Supplemental Info for "Evaluating the differential expression of TAM family receptors and efferocytosis activities in differentiated and polarized THP-1 macrophage"

**S1 Method. Image analysis using Fiji (ImageJ) Software.** Quantification of beads engulfed by macrophages was processed by Fiji, an open-source image analysis software (Schindelin, 2012). In brief, confocal images (6 slices of Z-stack images per treatment condition) of whole cell area stain (CP stain) (**Fig A**) and FITC labeled silica beads (**Fig B**) were analyzed for quantification. From Fiji, singular threshold values were chosen for each image to reveal only the whole cell area (**Fig C**) or the FITC beads area (**Fig D**). Threshold cell area was selected (**Fig E**) and quantified using the measurement tool within Fiji. The selected threshold cell area was then set as the "cell area mask" that was applied to the threshold bead area image (**Fig F**). This allowed for quantification of the bead area that was only contained within the cell, leaving out any stray, unengulfed beads that remained in the image wells after washing steps. The bead area was then quantified using the pixel area measurement tool. The two measurements were then used to calculate an engulfment score using **Eq. 1**.

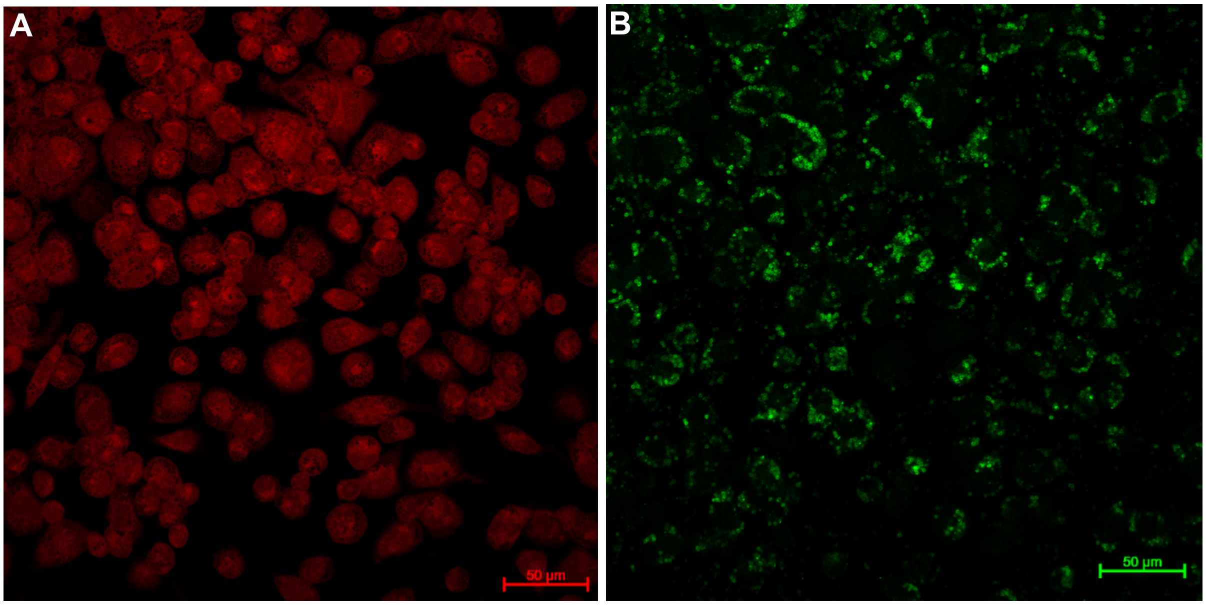


Representative confocal images of bead engulfment. **(A)** CP cell stain **(B)** FITC bead fluorescence.

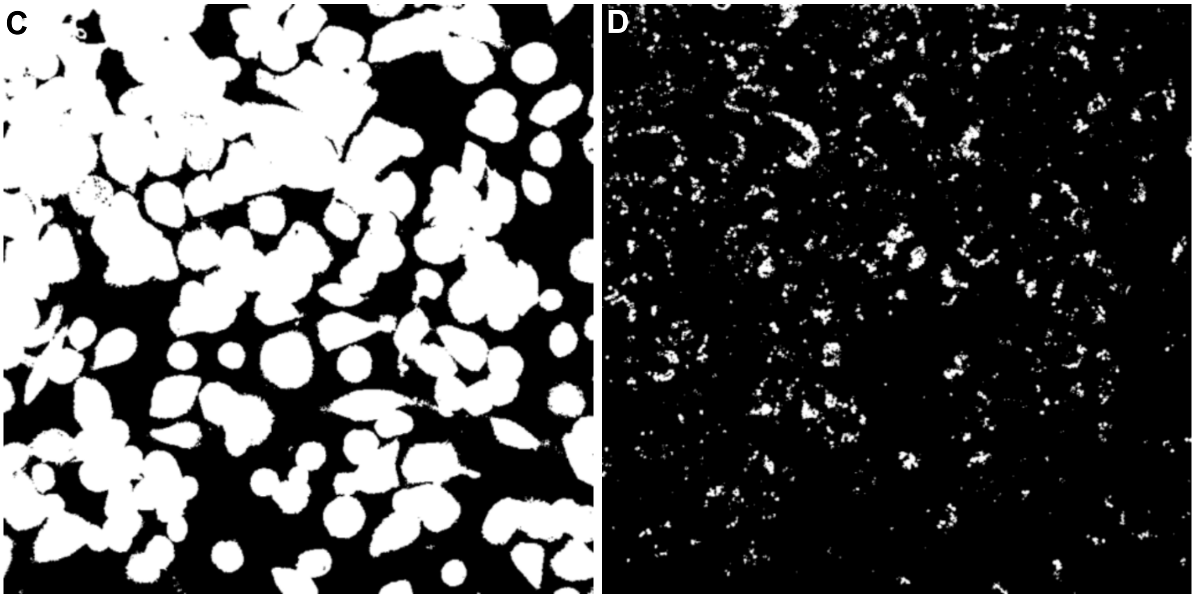


Threshold values set for both channels using ImageJ. **(C)** Cell stain by CP dye. **(D)** FITC bead fluorescence.

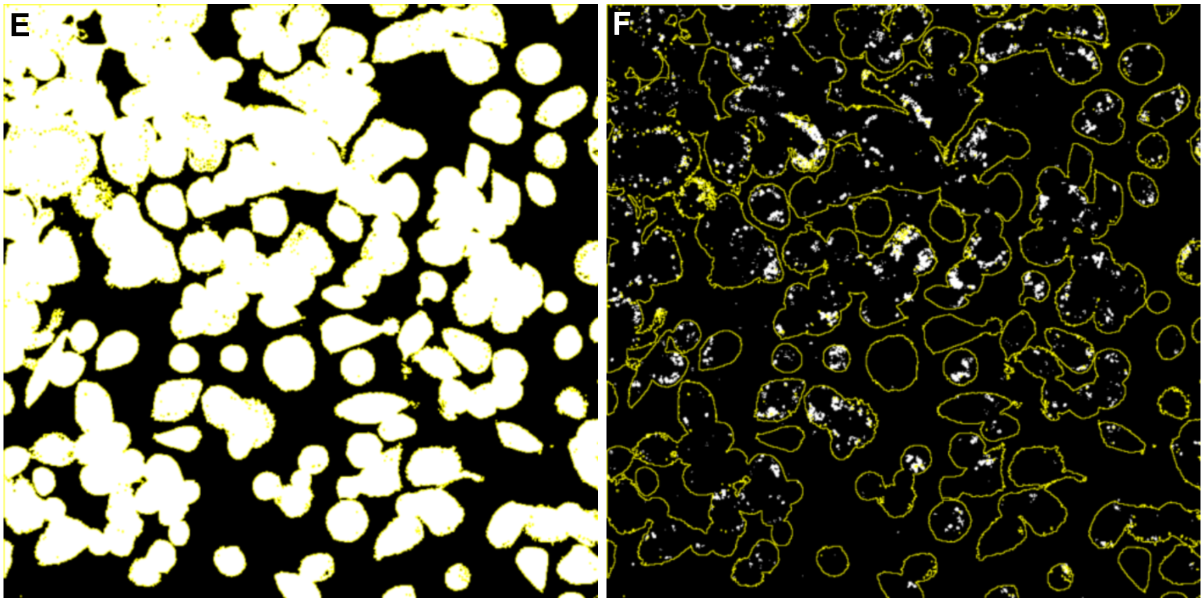


Selection of cell stain (CP dye) threshold file was made, and a measurement obtained. **(E)** Example measurement of pixels of CP stain. 70580.178 pixels measured in this example. **(F)** Once the cell stain pixel measurement is obtained, the cell stain selection was then transferred to the FITC channel to define the outline of individual cells. This border definition confines only the bead area within the cell, excluding any non-engulfed beads. In this example the beads channel showed 2121.192 pixels. The engulfment value of this example is therefore 3.005, calculated by the following equation:

$$\textbf{Eq 1}: Engulfment\ Value = \frac{Bead\ Area}{Cell\ Area} \times 100$$

$$Example: \frac{2121.192}{70580.178} \times 100 = 3.005$$

**S1 Figure. Bright field images of THP-1 derived M1 and M2 macrophages.** M$\phi$ cells were polarized into either the M1 and M2 macrophage cell types using IFN$\gamma$/LPS or IL4/IL13, respectively. Images were produced using Brightfield microscopy. M1 (Left), M2 (Right). Scale bar, 200 µm.

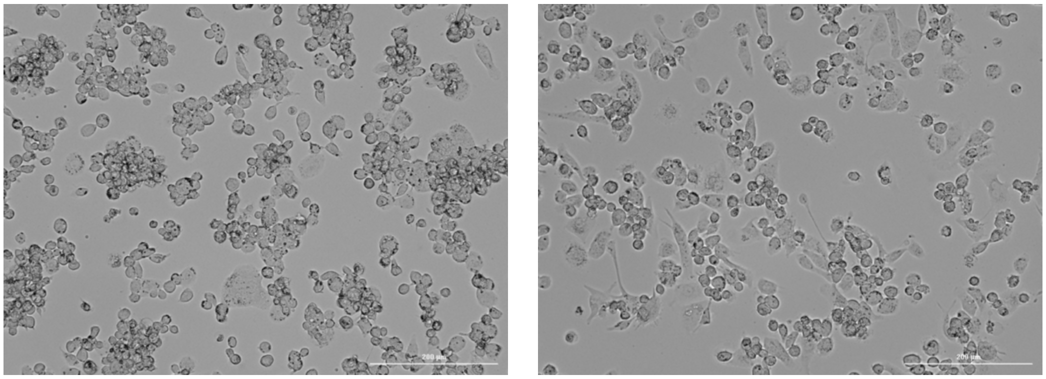

**S2 Figure. Minimal PROS expression in the THP-1 monocytes and derived macrophages.** THP-1 cells were differentiated using varied treatment conditions and were analyzed using western blot analysis. Recombinant Human PROS (20 ng) was used as a positive control.

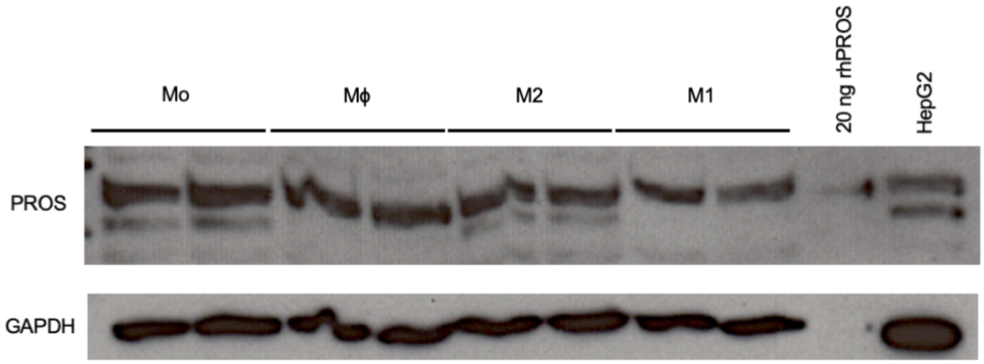

**S3 Figure. Expression patterns of TAM receptors in response to LPS, and Type I and II IFNs.** Western blot analysis of TAM receptors and GAS6 within THP-1 Mϕ stimulated with varying concentrations [above lanes] of LPS, IFNα, IFNβ and IFNɣ.

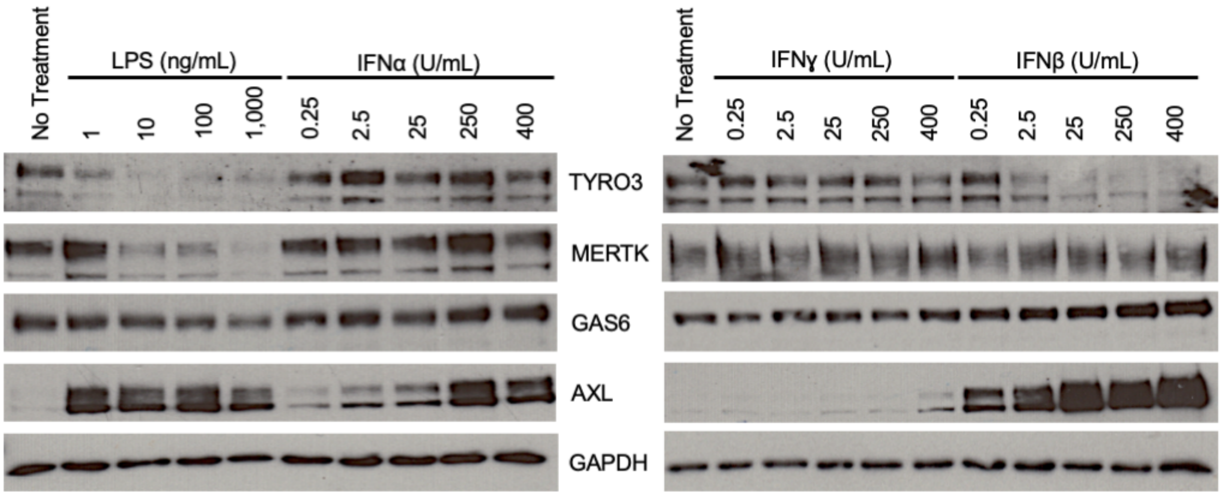

**S4 Figure. Annexin V saturation of PS-beads.** PS-beads were incubated with varying concentrations [0, 0.5, 0.736, 2.5, 5 µg/mL] of Annexin V-647 to visualize and quantify saturation of phosphatidylserine. Phosphatidylserine saturation was quantified by measuring the Annexin V-647 fluorescent intensity of each image.

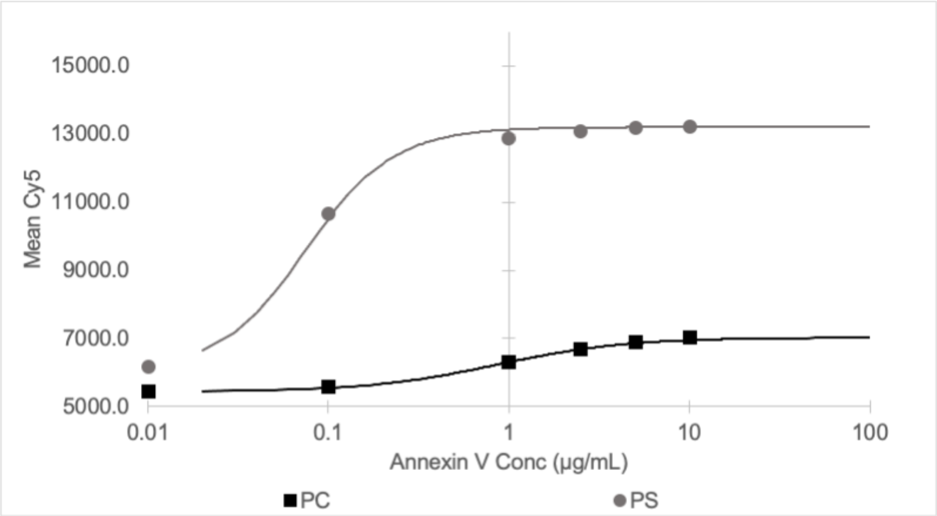

**S1 Table. Primary and Secondary antibodies used in the study.**

| Antibodies | Company | Cat# |
|---|---|---|
| Axl (C89E7) Rabbit mAb | Cell Signaling Technology | 8661 |
| CD11b/ITGAM (D6X1N) Rabbit mAb | Cell Signaling Technology | 49420 |
| CD14 (D7A2T) Rabbit mAb | Cell Signaling Technology | 56082 |
| Cleaved PARP (Asp214) (D64E10) XP Rabbit mAb | Cell Signaling Technology | 5625 |
| HA-Tag (C29F4) Rabbit mAb | Cell Signaling Technology | 3724 |
| MRC1/CD206 (C-10) Mouse mAb | Santa Cruz Biotechnology | sc-376232 |
| GAPDH (14C10) Rabbit mAb | Cell Signaling Technology | 2118 |
| Gas6 (D3A3G) Rabbit mAb | Cell Signaling Technology | 67202 |
| Mer (D21F11) XP® Rabbit mAb | Cell Signaling Technology | 4319 |
| Protein S (F-10) Mouse mAb | Santa Cruz Biotechnology | sc-271326 |
| Tyro3 (D38C6) Rabbit mAb | Cell Signaling Technology | 5585 |
| Pierce Goat Anti-Rabbit IgG, (H+L) Peroxidase Conjugated | ThermoScientific | 31460 |
| Pierce Goat Mouse IgG, (H+L) Peroxidase Conjugated | ThermoScientific | 31430 |

**S2 Table. Summary of AXL, MER, and GAS6 expression within the THP-1 macrophage cell-types.**

| | **Mo** | **Mϕ** | **M2** | **M1** |
|---|---|---|---|---|
| **AXL** | Hi | Low | Low | + |
| **MERTK** | - | + | Hi | Low |
| **GAS6** | Hi | Low | Hi | Low |
